# Mapping the signatures of inflammatory pain and its relief

**DOI:** 10.1101/2021.06.16.448689

**Authors:** Manon Bohic, Luke A. Pattison, Z. Anissa Jhumka, Heather Rossi, Joshua K. Thackray, Matthew Ricci, William Foster, Justin Arnold, Nahom Mossazghi, Max A. Tischfield, Eric A. Yttri, Ewan St. John Smith, Ishmail Abdus-Saboor, Victoria E. Abraira

## Abstract

Ongoing pain is often driven by direct activation of pain-sensing neurons and neuroimmune mediated sensitization. These heightened states of pain alter physiology, reduce motor function, and alter motivation to engage in normal behaviors. The complexity of the pain state has evaded a comprehensive definition, especially in nonverbal animals. Here in mice, we capture the physiological state of sensitized pain neurons at different time points post-inflammation and used computational tools to automatically map behavioral signatures of evoked and spontaneous displays of pain. First, retrograde labeling coupled with electrophysiology of neurons innervating the site of localized inflammation defined critical time points of pain sensitization. Next, we used high-speed videography combined with supervised and unsupervised machine learning tools and uncovered sensory-evoked defensive coping postures to pain. Using 3D pose analytics inspired by natural language processing, we identify movement sequences that correspond to robust representations of ongoing pain states. Surprisingly, with this analytical framework, we find that a commonly used anti-inflammatory painkiller does not return an animal’s behavior back to a pre-injury state. Together, these findings reveal the previously unidentified signatures of pain and analgesia at timescales when inflammation induces heightened pain states.

## Main

The sensitization of sensory neurons innervating injured tissue represents the first step in the transmission of pain^1^. Neuronal sensitization can trigger long lasting molecular and synaptic changes in peripheral and central circuits^2–4^. How these cellular changes align with behavior is central to understanding, quantifying and treating pain.

To first identify the time points that punctuate the development of injury-induced pain behaviors, we characterized the functional and molecular changes of sensory neurons that innervate injured tissue across time. We used the carrageenan model of localized pain because of its widespread use as an inflammatory pain model in rodents^5,6^. Carrageenan injection into the paw results in swelling of the whole paw and ankle at 4- and 24-hours post-injection^7^ (Fig. 1a, b, c). To determine changes to the excitability of isolated sensory neurons innervating the site of injury, we used the retrograde tracer Fast Blue^8,9^ (Fig. 1d). We found that ∼4.46% of cultured lumbar (L2 – L5) dorsal root ganglion (DRG) sensory neurons were Fast Blue positive (Extended Data Fig.1a) with cell body diameters resembling the natural distribution of sensory neurons that innervate the paw^10^.

**Figure 1.**
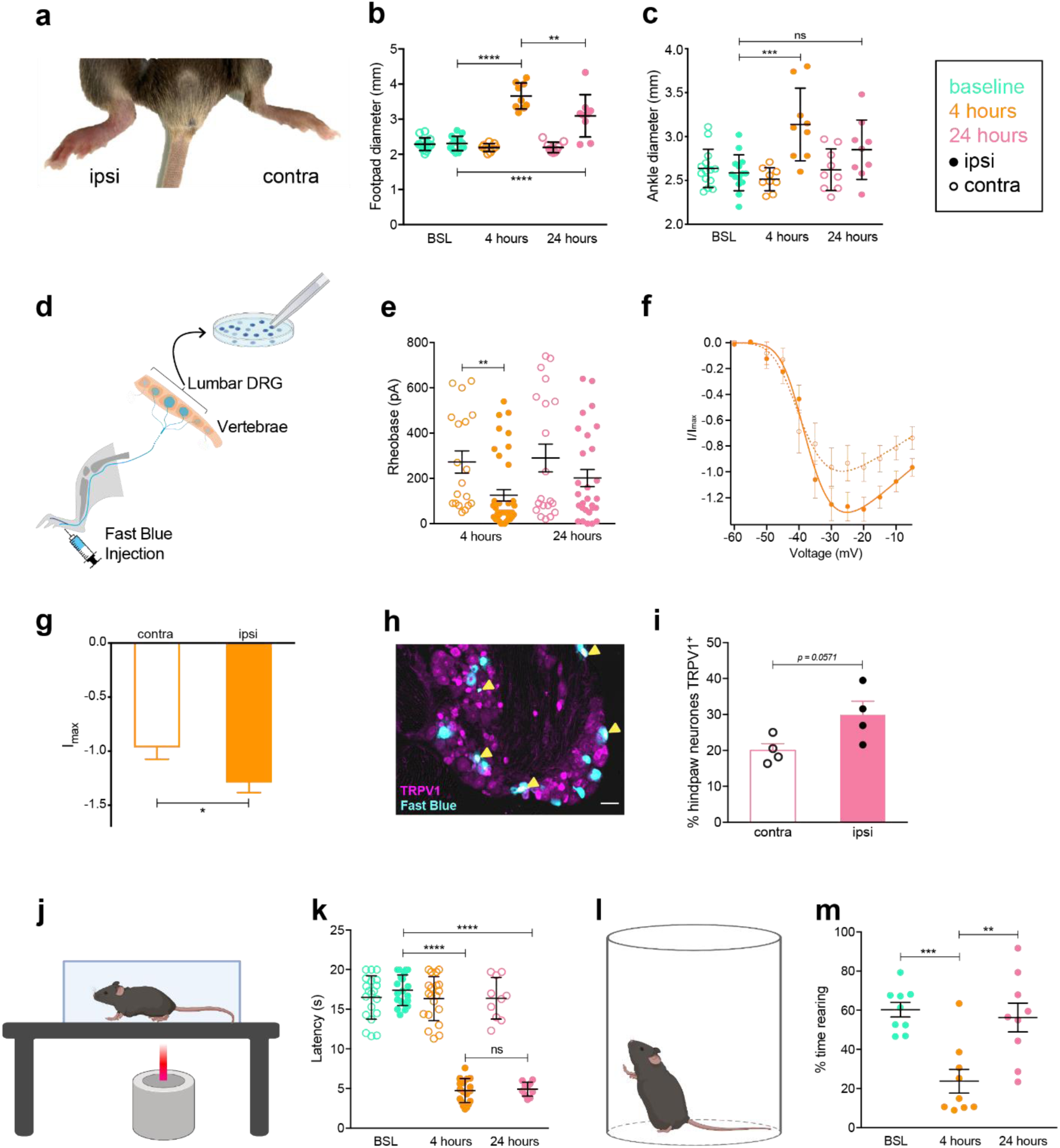
Unilateral injection of carrageenan into the hind paw of mice results in changes in nociceptor excitability and expression. **(a)** Noticeable inflammation of the injected (ipsi.) hind paw was observed 4-hours post-injection with carrageenan compared to the non-injected paw (contra.). Swelling of both the **(b)** footpad and **(c)** ankle was quantified for both hind paws with digital calipers before, 4-hours-post and 24-hours-post induction of inflammation with carrageenan. **(d)** Schematic representation of retrograde labelling of hind paw innervating sensory neurons with Fast Blue followed by cell culture and whole cell patch clamp electrophysiology, Insert: Fast Blue positive neuron (blue) following acute dissociation, scale = 50 µm. **(e)** Step-wise current injections were used to determine the rheobase of hind paw innervating sensory neurons from the ipsilateral and contralateral sides 4- or 24-hours post-induction of inflammation with carrageenan. Inward macroscopic current densities normalized to the average peak inward current density of contralateral cells **(f)** 4-hours post inflammation and **(g)** normalized peak inward current densities. **(h)** A subset of hind paw innervating sensory neurons (blue) express the nociceptive ion channel TRPV1 (magenta), cells positive for both Fast Blue and TRPV1 are identified by yellow pointers, scale = 50 µm. **(i)** A higher proportion of ipsilateral hind paw innervating neurons expressed TRPV1 after 24-hours of carrageenan-induced inflammation compared to the contralateral paw. **(j)** Schematic representation of mouse Hargreaves thermal pain assessment. **(k)** Hargreaves measurement of carrageenan-induced heat hypersensitivity at baseline and following 4- and 24-hours. **(l)** Schematic representation of traditional assessment of mouse spontaneous rearing behavior in a cylinder. **(m)** Time spent rearing by mice before, 4-hours-post and 24-hours-post induction of inflammation with carrageenan. * p < 0.05, ** p < 0.01, *** p < 0.001, **** p < 0.0001: **(b, c, k, m)** one-way ANOVA with Bonferroni post hoc; **(e, g)** unpaired t-test; **(i)** Mann-Whitney test.

Characterization of cultured neurons innervating the inflamed paw (ipsilateral) and non-injected paw (contralateral) revealed electrophysiological and molecular signatures that punctuate pain progression at 4- and 24-hours post-injury (Fig. 1d, e, f, g and Extended Data Fig. 1, Table1). Most notably, a lower rheobase of ipsilateral sensory neurons after 4-hours of inflammation (Fig. 1e), which suggests the rapid onset of peripheral sensitization correlating with the peak of physical inflammation at the paw (Fig. 1a, b, c). Indeed, voltage-gated inward currents, controlling the excitability of sensory neurons, show a significant increase in current magnitude at 4-hours, which is absent at 24-hours; no changes were observed in voltage-gated outward currents at any time point (Fig. 1 f, g, Extended Data Fig. 1b-l, Table1). At 24-hours, as inflammation subsides, and inflammatory mediators decline^11^, the degree of excitability of neurons also declines to match that of the contralateral side (Fig. 1e, Extended Data Fig. 1b-l). However, at 24-hours we see an upregulation of TRPV1 (Fig. 1h, i), an ion channel involved in the detection of noxious heat^12,13^. Thus, 4- and 24-hours post-inflammatory insult are critical time points to study the transition from acute changes in peripheral neuron excitability to longer-lasting molecular changes that underlie ongoing inflammatory pain.

**Table 1.** Electrophysiological characterization of cultured neurons innervating the inflamed paw reveal unique cellular signatures that punctuate the pain progression at 4- and 24-hours. Intrinsic and active properties of hind-paw innervating dorsal root ganglion neurons from the carrageenan injected side (Ipsi) and contralateral (Contra) side.

|  | Hours post-inflammation |  |  |  |
| --- | --- | --- | --- | --- |
|  | 4 hours |  | 24 hours |  |
|  | Contra (n = 20) | Ipsi (n = 39) | Contra (n = 20) | Ipsi (n = 28) |
| Resting membrane potential (mV) | -48.20 ± 1.49 | -46.05 ± 0.83 | -49.00 ± 1.43 | -45.64 ± 1.40 |
| Capacitance (pF) | 31.23 ± 2.69 | 25.90 ± 1.73 | 30.77 ± 4.44 | 33.17 ± 3.03 |
| Action potential amplitude (mV) | 80.33 ± 3.95 | 84.58 ± 2.17 | 85.45 ± 3.65 | 87.11 ± 2.74 |
| Half peak duration (ms) | 3.04 ± 0.26 | 3.50 ± 0.15 | 3.87 ± 0.35 | 3.37 ± 0.27 |
| Afterhyperpolarisation amplitude (mV) | 15.43 ± 1.30 | 14.10 ± 0.76 | 16.61 ± 0.71 | 16.15 ± 0.84 |
| Afterhyperpolarisation duration (ms) | 15.36 ± 1.55 | 17.82 ± 1.55 | 15.48 ± 1.17 | 16.45 ± 1.19 |

Having established the critical time points for inflammatory pain progression, we next used traditional measures of evoked pain behavior to differentiate 4- vs. 24-hours pain states. We measured the latency of paw withdrawal to heat with Hargreaves, one of the most commonly used assays to measure noxious heat hypersensitivity following inflammation^14^. This showed strong hypersensitivity at both 4- and 24-hours, with no distinguishable change between the two time points (Fig. 1j, k). By contrast, measuring ethological behaviors, such as time spent rearing, can better resolve these two time points (Fig. 1l, m). Taken together our results demonstrate that while key biological changes to sensory neurons punctuate pain progression over time, current binary algesiometric assays for sensory evoked behavior do not have the proper dynamic range to resolve the time course of injury and healing.

### High-speed videography of sensory-evoked reflexes resolve injury progression and differentiate allodynia from hyperalgesia

To increase the dynamic range of sensory-evoked behavior assays we used high-speed videography to break down paw withdrawal to sensory stimuli into sub-second movements within groups of recently established short-latency reflexive vs. long-latency affective behavioral families^15,16^ (Fig.2a-c). Mechanical hypersensitivity is a common complaint of inflammatory pain, presenting itself as either allodynia, when innocuous sensations become painful^17–19^, or as hyperalgesia, when there is an increased sensitivity to noxious sensations^20,21^. While these two conditions can easily be assessed in the clinic^22,23^, differentiating them in non-verbal animal models is a challenge^24,25^. Thus, with high-speed videography we recorded the animal’s response to both innocuous (brush) and noxious (pinprick) stimulations. With this strategy we did not observe a significant difference between baseline, 4- and 24-hours in the short-latency reflexive behavioral families to either innocuous (brush) or noxious (pinprick) stimulations of the paw (Extended Data Fig.2 a, b). However, coping behaviors (paw shaking and paw guarding) associated with affective behavior families, are more dynamically regulated at 4- vs. 24-hours post-injury in response to brush and pinprick (Fig. 2d, e). Indeed, while paw guarding duration evoked by brush is significantly increased at both 4- and 24-hour time points, paw guarding duration evoked by pinprick is only upregulated at 24-hours (Fig. 2d, e).

**Figure 2.**
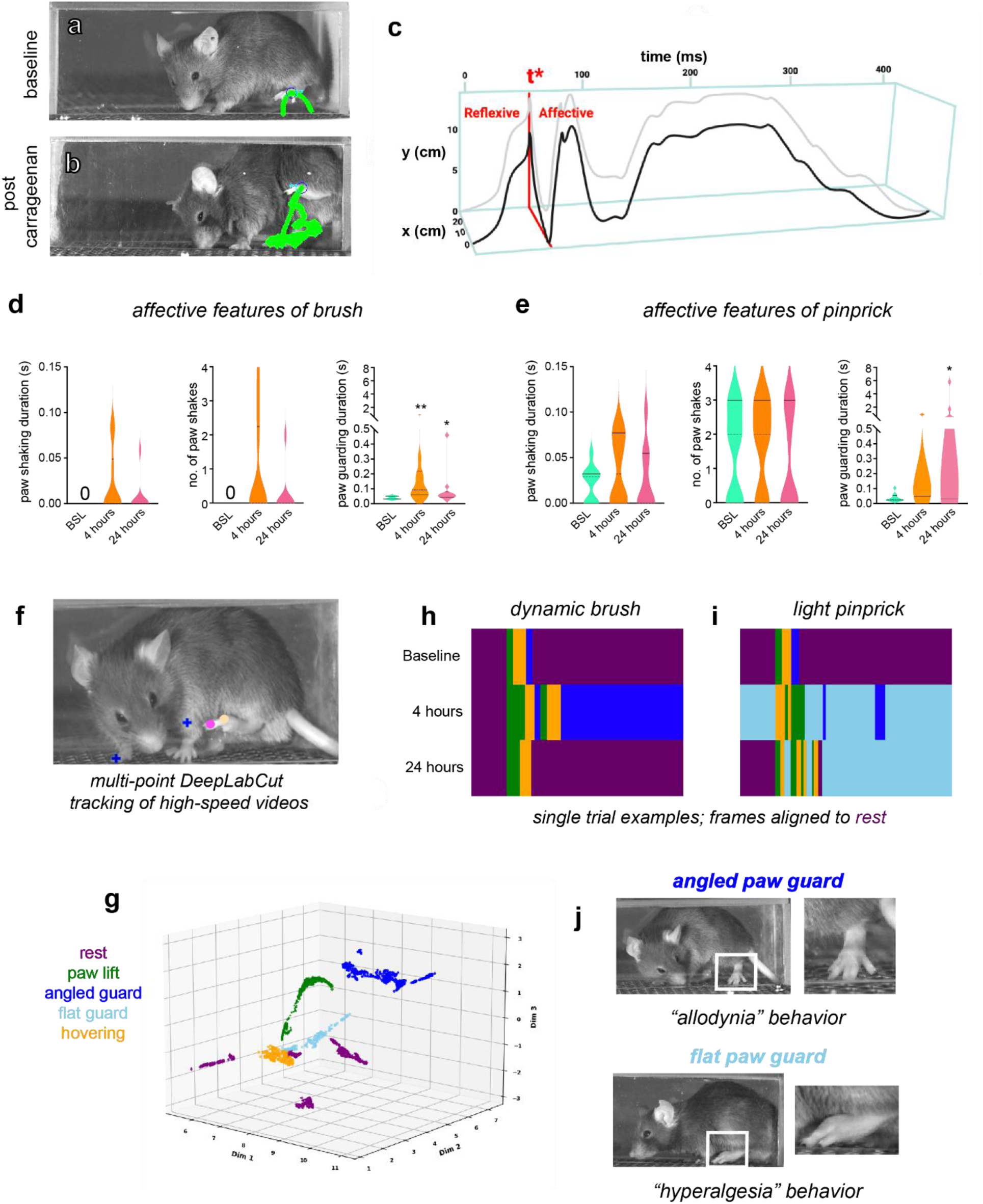
PAWS and B-SOiD automated pain assessment platforms detect defensive coping behaviors associated with pain sensation during inflammation. **(a)** A behavioral response to a somatosensory stimulus at baseline. **(b)** Post-carrageenan injection mice guard the paw in the air for extended time. **(a,b)** Green lines show paw trajectory pattern across entire behavior, and mouse image shows single frame with paw at its apex. **(c)** Image drawn in Biorender and modified from Jones et al., 2020. PAWS software measures paw height, velocity, shaking, and guarding in X and Y axes. Reflexive behaviors occur before paw reaches its apex (t*), whereas affective behaviors occur after t*. **(d,e)** Affective features such as paw guarding are upregulated in response to dynamic brush and light pinprick comparing baseline to 4- and 24-hours post-carrageenan injection. **(f)** Identification of inputs to be used for unsupervised classification in B-SOiD. **(g)** Low-dimensional projection of identified feature clusters. Colors of sub-clusters indicate their post-hoc behavioral group assignment. **(h)** Representative responses to dynamic brush and **(I)** light pinprick at baseline, 4-hours, and 24-hours time points post-carrageenan injection. Responses are color-coded by the identified action type as in panel **(g). (j)** Examples of the two forms of guarding identified by B-SOiD, which may be indicative of the activation of different subsets of sensory neurons (mechanoreceptors by brush, inducing angled guard, nociceptors by pinprick, inducing flat guard). N=10 mice per group and Wilcoxon matched-pairs sign rank test was performed to determine statistical significance. * p < 0.05, ** p < 0.01

To further differentiate the long-latency affective behavioral signatures driven at 4- vs. 24-hours post-injury, including signatures specific to noxious and innocuous stimuli^26,27^, we employed a recently developed unsupervised machine learning approach to parse spatiotemporal patterns in paw position data (B-SOiD^28^). For accurate comparison, we used the same high speed behavioral data collected in Fig. 2a-e. As inputs, we used two positions within the hind paw and two reference points as identified with the deep neural network DeepLabCut (Fig. 2f, see Methods). B-SOiD then identified and extracted unique clusters of conserved motor responses to these stimuli (Fig. 2g; categorical names were assigned to clusters post hoc). We found eleven sub-action clusters across stimulation contexts. Compared to baseline, we observed changes in response to stimuli at 4- and 24-hours post-injection (Fig. 2h-j). Notably, B-SOiD detected increased guarding across stimuli. Additionally, a lift-to-hover pattern was identified as a behavioral module distinct from guarding. This lift-to-hover pattern was common at baseline, but with a brush stimulus became extended over time (Fig. 2h). These behaviors and distributions are similar to those observed by our top-down supervised approach (Fig. 2d, e), with notable distinctions. B-SOiD extracted two guarding types specific to the foot stimulus presented (i.e. angled versus flat guard) (Fig. 2j). These phenotypes were distinct from each other in both height and foot posture. The upregulation of the “angled guard” (characterized post-hoc as paw lifted and perpendicular to the surface) with the innocuous stimulus, suggests a behavioral response specific to mechanical allodynia^29^. Conversely, the “flat guard” was upregulated with the noxious stimulus (paw lifted and parallel to the surface), likely associated specifically hyperalgesia^30^ (Fig. 2j).

Altogether, our approach can finely measure the transition in inflammatory-induced sensitization at 4- and 24-hours and highlights the importance of affective behavioral biomarkers as representations of inflammatory pain. We find that inflammatory pain specifically alters the responses to mechanical stimuli such that defensive coping behaviors are more frequent. Interestingly, we show that mechanical allodynia (e.g. hypersensitivity to innocuous stimuli) appears as early as 4-hours while mechanical hyperalgesia (e.g. hypersensitivity to painful stimuli) is most upregulated at 24-hours. To our knowledge this is the first evidence of a robust and generalizable sensory-evoked behavior feature that distinguishes mechanical allodynia versus mechanical hyperalgesia in rodents.

### Ethological approach to movement-evoked spontaneous measures of inflammatory pain

Sensory-evoked responses alone do not accurately reflect the most common symptoms experienced by chronic pain patients^31^. Movement-evoked pain is significantly greater and a far more common clinical problem than sensory hypersensitivities^32^. Our findings that ethological behaviors can better define pain progression over time (Fig.1k) prompted us to explore unbiased approaches to scoring behaviors in freely moving mice at 4- vs. 24-hours post-injury. To detect, measure and scale behavior in freely moving animals, we used time-of-flight infrared cameras to detect mouse body contours, depth and movement during 20-min long sessions (Extended Data Fig. 3). We then applied 3D pose analysis using unsupervised machine learning^33,34^ to identify sets of sub-seconds long movements (a.k.a. “modules”) that best categorize families of behavior at 4- vs. 24-hours post-injury (Fig. 3a, b, c). To identify ethologically meaningful modules representative of spontaneous and movement-evoked pain, we classified the 62 identified modules as belonging to one of four types of behavior: exploration, grooming, pausing and rearing (Table2). We found that all four types of behaviors were affected as a result of inflammation (Table2). In accordance with traditional measures showing a transient decrease in rearing behavior at 4-hours (Fig. 1k), we found that the usage of rearing modules is overall downregulated at 4-hours (module 42, Fig. 3d). However, we found a prolonged impact of pain on most rearing behaviors at both 4-hours and 24-hours. This result suggests that rearing actually remains painful for animals at 24-hours, highlighting ongoing pain often observed following inflammation and sometimes missed by traditional approaches. Interestingly, we found that grooming was mainly downregulated by ongoing pain (module 43, Fig. 3e) while pausing modules significantly affected in pain states were mostly upregulated (module 50, Fig. 3f). Thus, inflammatory pain induces a mosaic of behavioral changes that we can resolve with 3D pose estimation analysis. Consistent with this observation, Linear Discriminant Analysis (LDA) of module usage across time allows us to visualize globally the transformation of spontaneous behavior to adapt to a new pain state (Fig. 3c). The progressive transformation we uncover here resolves the discrepancy between the punctual change in observable anatomical parameters of inflammation, and in rearing behavior as measured by traditional approaches at 4-hours (Fig. 1b, c, k) and the ongoing evoked pain at 4- and 24-hours (Fig. 1i).

**Figure 3.**
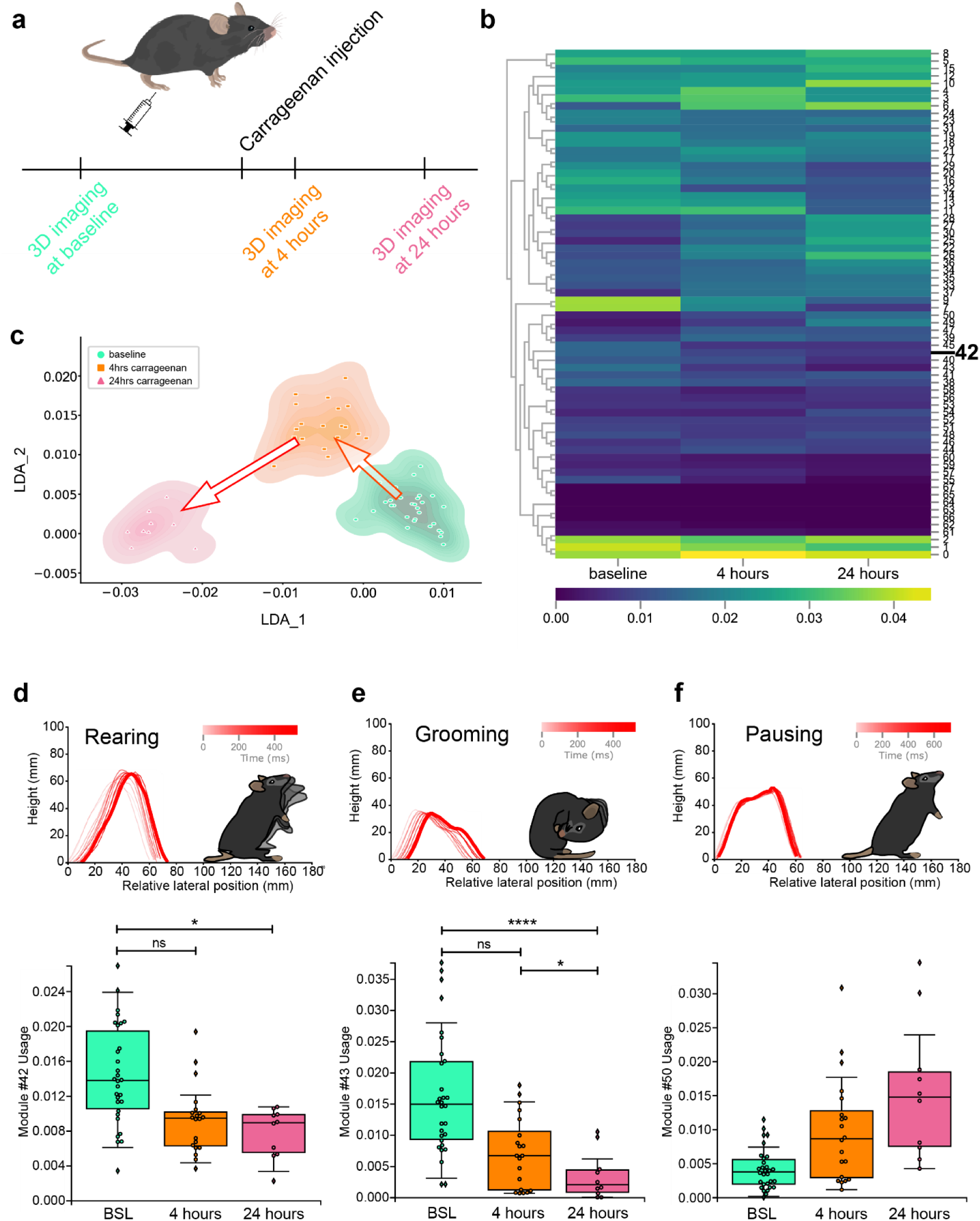
3D pose analysis detects behavioral signatures of carrageenan-induced inflammatory pain. **(a)** Timeline of the experiment. **(b)** Heatmap representation of changes in module usage by the three experimental animal groups baseline, 4-hours and 24-hours post-carrageenan injection. **(c)** Representation of linear discriminant analysis (LDA) of spontaneous behavior module usage at baseline, and following carrageenan injection at 4-hours, 24-hours. **(d,e,f)** Example of modules with varying usages at baseline, and following carrageenan injection at 4-hours, 24-hours with a spinogram in red, mouse representation of the micro-movement and usage. **(d)** Example of a rearing module (42, rear against the wall) less used at 4- and 24-hours. **(e)** Example of a grooming module (43) less used at 4- and 24-hours with a significant difference between 4- and 24-hours behavior. **(f)** Example of a pause module (50, pause with head up/observe) more used at 4- and 24-hours.

**Table 2.**
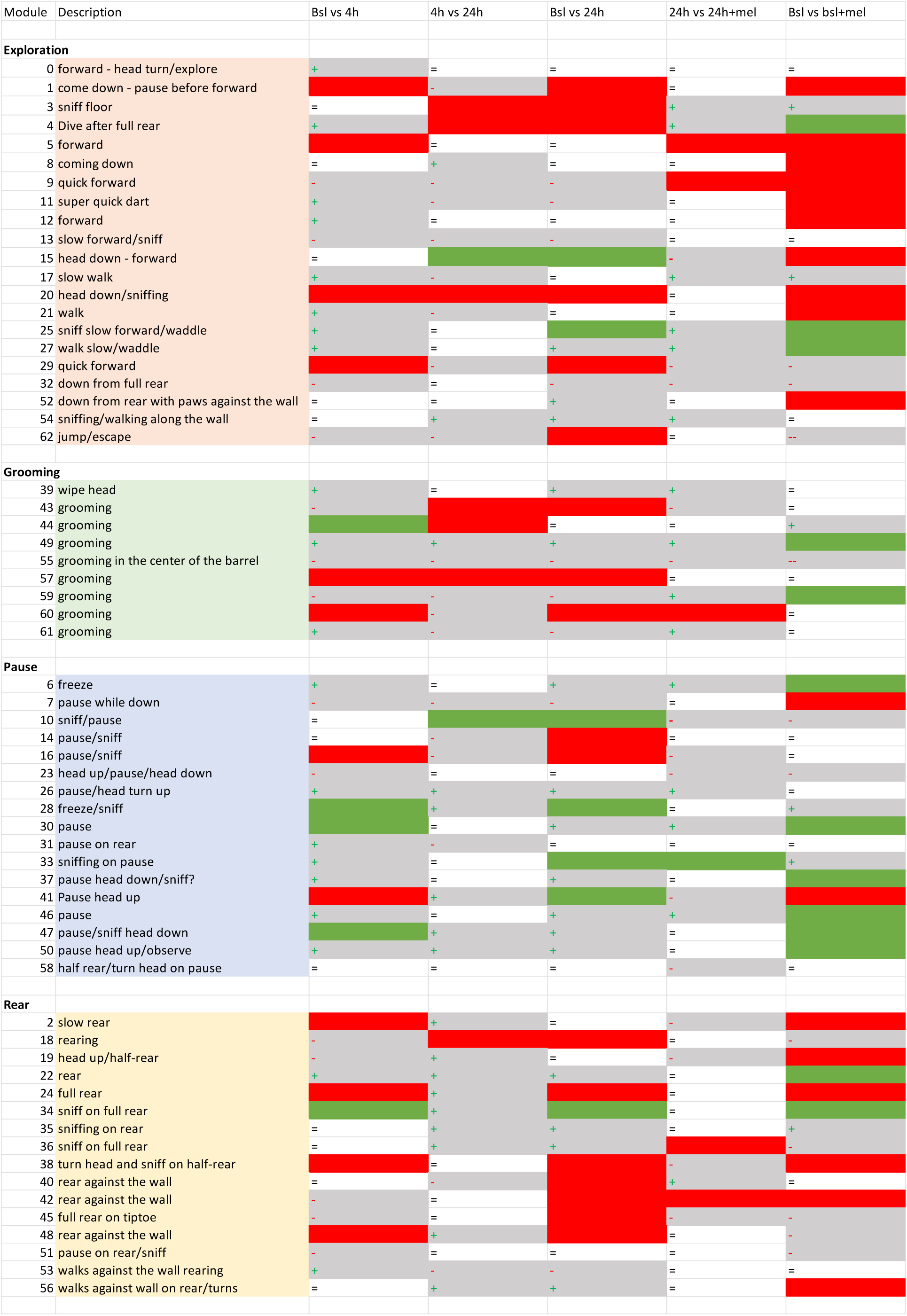
Mouse spontaneous behavior modules pre and post-injury, and in analgesic state. Annotation and classification of the set of 62 modules identified by 3D pose analysis as representative of mouse spontaneous behavior pre-injury, at 4- and 24-hours post-carrageenan injection, 24-hours post-injection with meloxicam IP injection and pre-injury with meloxicam IP injection. Modules in red were less used, modules in green were more used (statistically significant bootstrap t-test).

Altogether, we find that 9 out of 62 modules are differentially regulated at 4- vs 24-hours post-injury. We propose that these modules represent a cohort of sensitive behavioral biomarkers of inflammatory pain, central to understanding the neuronal networks driving pain progression over time. For example, resolving precisely how these specific spontaneous pain signatures correlate to neuronal activity driving plasticity changes from peripheral to central mechanisms is crucial to targeted analgesic development.

### Meloxicam relieves hyperalgesia but does not return behavior to a pre-injury state

To validate both sensory-reflexive (Fig.2) and spontaneous (Fig.3) inflammatory pain behavioral biomarkers, we explored the effects of the anti-inflammatory drug meloxicam on the usage of these biomarkers at 24-hours (Fig. 4a). Meloxicam is commonly used to relieve pain and inflammation in rodents, dogs, cats and humans^35–42^. At 22-hours post-carrageenan paw injection, we injected mice with either meloxicam or saline intraperitoneally and assessed heat hypersensitivity with Hargreaves, sensory-reflexive responses with high-speed videography, and spontaneous behaviors with 3D pose dynamic analysis (Fig.4a). As expected, we found a reduction in inflammation-induced heat hypersensitivity (Fig.4b). We then tested sensory-reflexive responses with high-speed videography and machine learning (Fig.4c-d). Similar to our observations above (Fig.2d-e), after inflammation, we observed significantly upregulated paw guarding duration evoked by 1) dynamic brush stimulation at 4-hours (Fig. 4c) and 2) light pinprick stimulation at 4- and 24-hours post-carrageenan injection (Fig. 4d). Interestingly, meloxicam administration prevented the progression of carrageenan-induced paw guarding duration in response to light pinprick typically observed at 24-hours (Fig. 4d). Combined with the hyperexcitability of injury-innervating neurons we observed at 4- but not 24-hours (Fig. 1e), this suggests that prolonged inflammation results in sensitization of central circuits that drive tactile hyperalgesia and meloxicam can target this secondary effect of inflammation to blunt this behavior.

**Figure 4.**
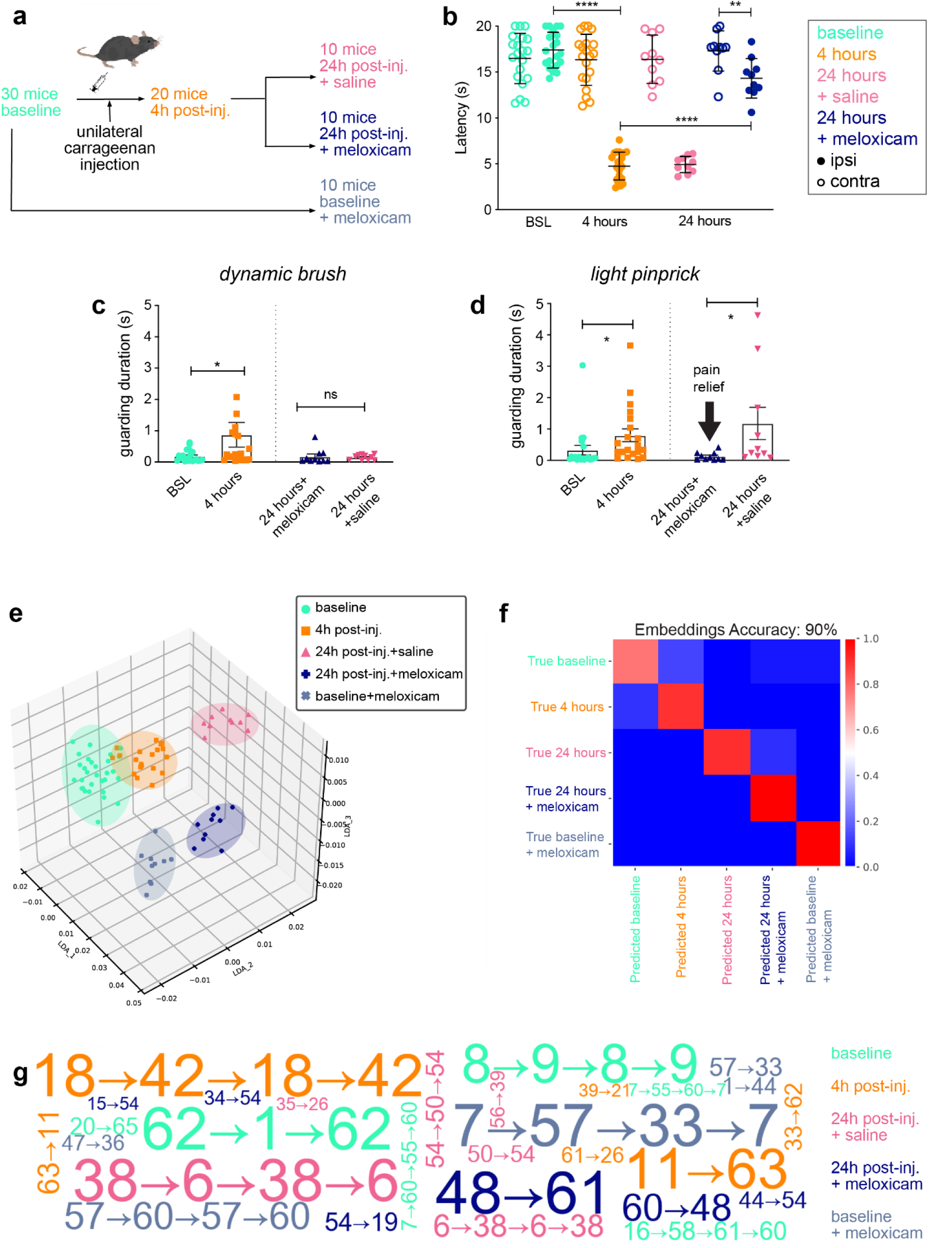
Machine learning and natural language processing approaches show that meloxicam relieves affective features of hyperalgesia but it does not promote return to pre-inflammation spontaneous behavior. **(a)** Timeline of the experiment. An initial group of 30 mice is tested at baseline with MoSeq and Hargreaves. A subgroup of 20 mice receives an intraplantar injection of 20 µl 3% carrageenan, is tested at 4h post-injection with MoSeq and Hargreaves, then further subdivided into two groups of 10 mice injected intraperitoneally with saline or meloxicam respectively at 22h and again tested with MoSeq and Hargreaves at 24h. A control group of 10 mice is injected with meloxicam at baseline and tested with MoSeq and Hargreaves. **(b)** Hargreaves measurement of carrageenan-induced heat hypersensitivity at baseline, 6h following carrageenan injection, as well as pain relief by meloxicam at 26hrs compared to saline. **(c,d)** Paw guarding duration is measured at baseline, and 4h post-carrageenan injection, and 24h post carrageenan injection after saline or meloxicam intraperitoneal injection (striped and solid bar respectively) following dynamic brush **(c)** or light pinprick **(d)**. **(e)** 3D pose analysis of 5 groups: baseline, baseline + meloxicam intraperitoneal injection, 4h post-carrageenan injection, 24h post-carrageenan injection + saline intraperitoneal injection, 24h post-carrageenan injection + meloxicam intraperitoneal injection. **(e)** Representation of linear discriminant analysis (LDA) of raw usage for the five different groups. **(f)** Confusion matrix for a classifier trained on learned embedding with leave-one-out cross validation. Accuracy of prediction of the different experimental groups reaches 90% with a context-dependent learned embeddings approach inspired by natural language processing. **(g)** Word cloud representation of the top most representative 2-, 3- or 4-long module sequences for each experimental group as extracted with the learned embeddings method. A standard co-location algorithm was first used to detect 2-long module sequences according to whether the 2-long sequence (e.g. A→B) appeared significantly more than each of its constituents (i.e. A or B). Wherever a significant 2-long sequence was detected, we replaced it by a new agglomerated syllable representing the co-location. We then recursed on this procedure to find 3- and 4-long sequences, at each iteration checking for the significance of agglomerated sequences by comparing their frequency to those of each of the sequence’s constituents. Briefly, we found that escape behavior is most representative of mice at baseline (62→1→62, jump against the wall→down→jump against the wall). A specific sequence of rearing best describes mouse spontaneous behavior in the early onset of inflammatory pain (18→42→18→42, rear against the wall→rear and sniffing) which could be interpreted as an adaptation of the escape behavior observed at baseline to a new pain state. After 24 hours in pain, a more subdued type of exploratory behavior (38→6→38→6, turn head and sniff on half-rear→freeze) is most representative of ongoing inflammatory pain. Finally, while the anti-inflammatory drug meloxicam significantly relieves mechanical and thermal evoked pain at 24-hours, we find that a full rear against the wall followed by grooming is the behavioral sequence most representative of spontaneous behavior after administration of meloxicam to mice in pain (48→61). Interestingly, we also find a complex grooming sequence to be most representative of spontaneous behavior when meloxicam is administered at baseline in the absence of injury (7→57→33→7, pause→grooming→sniffing→pause), which suggests that a change in grooming behavior could represent a side-effect of meloxicam at baseline and in a pain state.

Next, we assessed the effect of meloxicam on identified movement-evoked pain signatures (Fig. 3). We found that meloxicam reinforced the shift in spontaneous behavior we observed in animals in pain, i.e. even less rearing, grooming and even more pausing (Extended data Fig. 4a, b, c). Consistent with individual module results, LDA shows that spontaneous behavior after meloxicam injection maps onto a very different pose space compared to baseline, 4-hours and 24-hours with saline injection (Fig. 4e). Interestingly, the spontaneous behavior of animals in pain that received meloxicam (Fig. 4e, dark blue) was strikingly different from that of animals pre-injury (Fig. 4e, green). That is, conversely to sensory evoked behaviors, meloxicam does not seem to alleviate spontaneous pain (Fig. 4e). This unexpected finding led us to question whether meloxicam might alter baseline behavior in the absence of inflammatory pain. Our analysis demonstrates that meloxicam does indeed affect spontaneous behavior even when administered in uninjured animals (Fig. 4e). Furthermore, we see that a meloxicam induced analgesic state (Fig.4e, dark blue) maps much closer in pose space to uninjured animals that received meloxicam (Fig.4e, light blue).

### Higher order behavioral sequences predict pain and analgesic states in rodents

While sub-second movement-evoked pain signatures can be resolved with module analysis (Fig.3), ongoing pain signatures necessitate a different analytical approach, one that can describe the structure of behavior over a longer time scale. Intuitively, we know that an injured knee can change the way we walk, abdominal pain can change the way we stand, and chronic pain can change the way we interact with our environment and with each other. To quantitatively represent these types of behavioral sequences, we calculated transition probabilities by counting the total number of occurrences where module A is followed by module B, for all modules. To first test whether analysis of transition probabilities can be used to describe ongoing pain signatures, we used a Sankey diagram to represent transition probabilities between incoming and outgoing modules based on root modules whose usage changes over time: 42 (rear against the wall) and 43 (grooming behavior, Extended Data Fig. 4d, e, f and g, h, i). We find that most sequences are stable across time, for example, the probability that 42 is preceded by 18 and followed by 1 (blue, Extended Data Fig. 4d, e, f). But a small number of sequences appear in pain states: 43→6 and 43→11 at 4-hours or 43→49 at 24-hours (pink, Extended Data Fig. 4g, h, i). We hypothesized that these unique novel sequences could be representative and discriminative of various pain states (4- vs 24-hours).

If we compare the analysis of spontaneous behavior to deciphering a new language, module usage informs us on word frequency, which holds only limited meaning as to the state of the animal. However, extracting sequences of modules, akin to deciphering the meaning of entire sentences^43,44^, might provide a better representation of ongoing pain states. To test this model, we applied standard sequence classification techniques of natural language processing^45^ to extract sets of modules which best represent a particular experimental group (baseline, 24-hour post-injection + saline, or 24-hour post-injection + meloxicam pain relief, for example). These methods embed long sequences (i.e., the raw sequencing data produced by one animal) in a representational space where sequences having similar co-occurrence structure (pairs of modules, triples of modules, etc.) tend to cluster. Consequently, these embeddings depend on contextual information^46^ which is absent in module usage data and potentially more powerful than first order transition probabilities (see Materials and Methods, Learned Embeddings of Module Sequences). To evaluate the relative predictive powers of usages versus first order transition probabilities versus learned embeddings, we trained three classifiers to predict experimental groups from each type of representation (see Materials and Methods, Classifier Analysis of Animal Representations). Context-dependent, learned embeddings (Fig.4f) were substantially more predictive of experimental groups than raw usages or transition probabilities (71.25 % accuracy for raw usages; 81.25 % accuracy for transition probabilities; 90.00 % accuracy for learned embeddings; full confusion matrices depicted in Extended Data Fig. 4j).

Learned embeddings now provide us with the means to identify precise behavioral sequences that characterize different pain states. To apply this finding, we adapted standard module co-location metrics^46^ to detect 2-, 3- and 4-long modules that characterized each of the five experimental conditions (Fig. 4g). These methods proceed by recursively agglomerating neighboring modules into higher-order units according to whether the higher-order unit appears significantly more than its constituents in the full sequence. This allowed us to identify sequences of spontaneous behavior most representative of various mouse internal states such as acute pain, adaptation to pain, effect of an anti-inflammatory drug on ongoing pain as well as its potential side-effects pre-injury. Mainly we found that sequences [62→1→62], [18→42→18→42], [38→6→38->6], and [48→61] best represent baseline, 4-, 24-hours post-injury, and 24-hours post-injury with meloxicam respectively (Fig. 4g). Ethological evaluation of these sequences (Table 2) suggests a shift in combinations of rears, jumps, sniffs, freezes, and grooming bouts that best represent acute, ongoing pain, and an analgesic state. Interestingly, we find unique sequences of grooming as the most representative spontaneous behavior after meloxicam injection both at baseline and in a pain state ([48→61] and [7→57→33→7]).

We then ablated these specific behavior sequences (constituting only 0.732 % of the total modules) by replacing them with random modules to re-learn embeddings on these ablated sequences. Despite removing only a small portion of the total modules, this procedure resulted in an average drop in classifier accuracy of 6.76 % +/- .017 % (s.e.), placing it within a standard deviation from the performance of the transition representation (Extended Data Fig. 4k). In contrast, when the same number of random modules were ablated, accuracy only dropped an average of 1.63 +/- 3.58 % (s.e.) (Extended Data Fig. 4k). We take this quantitative result as a proof of concept not only that complex behaviors beyond usages and transitions characterize different pain states but also that these complex behaviors can be detected using standard sequence representation methods from machine learning.

In conclusion, we have identified the learned embeddings method as a crucial tool in order to extract biologically meaningful data from rich and complex behavior datasets. This allowed us to uncover specific behavioral signatures of spontaneous pain as well as quantify the efficacy and side-effects of the anti-inflammatory analgesics.

## Discussion

We provide a holistic assessment of inflammatory pain behavior in the mouse that is correlated to key changes in neuronal excitability. With videography across timescales followed by unbiased analyses using machine learning, we identify evoked and spontaneous behaviors of inflammatory pain previously undetected with traditional methods. Profiling sensory neurons directly innervating the site of inflammation revealed that changes in neuronal excitability initiate inflammatory pain, but that persistent pain is driven by molecular changes^1,47,48^. We determined that meloxicam mediated pain relief does not equal a full return to a pre-inflammatory state. Taken together, these data provide a new multidisciplinary approach for arraying the high dimensional pain behavior datasets and offer a new experimental roadmap for assaying analgesic efficacy in preclinical rodent models.

### Resolving pain behaviors across time

A single injection of carrageenan produces localized inflammation, which starts resolving within a day, but pain-like behavior can persist for days. We demonstrate higher excitability of sensory neurons innervating the inflamed hind paw after 4-hours of inflammation. This rapid onset of hyperexcitability, as well as the pain relief mediated by meloxicam, point to mechanisms such as the release of prostaglandin E2 (PGE2)^11,14^. However, our multi-method approach shows that sensory neurons innervating the site of inflammation are no longer hyperexcitable 24-hours after inflammation. This suggests the persistent pain behaviors that we resolve at the 24-hour time point are likely driven by adaptations in the entire nociceptive system. Indeed, we show an increase in the number of sensory neurons expressing TRPV1 24-hours after induction of inflammation. Our results show that the status of sensory neurons evolves rapidly with inflammation, with unique changes in the activity and molecular profile of sensory neurons that are punctuated at discrete time points post-injury. This is consistent with recent findings from the RNA profiling of sensory neurons in a longer-lasting inflammatory pain model (Complete Freund’s Adjuvant (CFA) paw injection) showing an increase at 48-hours in PGE2 synthase expression in peptidergic nociceptors and TRPV1 in peptidergic and non-peptidergic nociceptors^49,50^. Developing, screening and testing single and combination therapies to treat pain over time requires an ability to differentiate between these critical time points with unique and easily identifiable behavior signatures.

### Supervised and unsupervised learning approaches to scale sensory-reflexive paw withdrawal behaviors can distinguish hyperalgesia and allodynia

Using the 4- and 24-hour time points to anchor our sensory-reflexive behavioral studies we show that the early onset of paw withdrawal behaviors are unaltered over time (paw speed and height, Fig.2). However, we observe an upregulation of defensive coping behaviors like paw guarding as pain progresses from 4- to 24-hours. This is consistent with recent work demonstrating that supraspinal brain structures like the parabrachial nucleus of the brainstem and the central and basolateral amygdala coordinate more complex defensive coping behaviors, like attendance to the stimulated area and escape behavior, at longer time scales^51–54^. Taken together, our results show a distinction between strictly reflexive and supraspinal mediated coping behavioral responses to sensory stimulation under pathophysiological conditions, that can be distinguished across time and are specific to different sensory modalities. In addition, our work shows identifiable variations in guarding responses to innocuous and noxious stimuli. To our knowledge, this is the first demonstration that differences in paw guarding behaviors are specific to a given mechanosensory modality, finally allowing for the nuanced distinction between allodynia and hyperalgesia in preclinical animal models.

### 3D pose estimation establishes spontaneous pain signatures and challenges the definition of analgesia in preclinical rodent models

While evoked measurements are necessary for the estimation of rodent pain^55^, our addition of 3D pose estimation to scale spontaneous pain behaviors provides a more rounded and unbiased picture of the overall rodent pain state, which is a much more accurate representation of the human pain experience. Our 3D pose estimation studies uncover two distinct trends in how ongoing pain affects general behavior. First, we show that mice differentially use unique micro-movements at 4-hours and even more so at 24-hours compared to pre-injury, highlighting a heightened pain experience at 24-hours. Second, we find a parallel trend that follows a U-shaped curve where the usage of certain movements is most affected during the acute phase and tends to revert to pre-injury levels at 24-hours. This second trend suggests an adaptation of motor behavior to pain.

The International Association for the Study of Pain describes pain as both a sensory and an emotional experience^56^, with analgesia broadly defined as a lack of pain or an insensitivity to pain. By this definition an analgesic would provide relief from both sensory-evoked and ongoing emotional pain. But is analgesia a return to a pre-injury state or a different state altogether? Our combined approach shows that each pain state (pre-injury, 4-, 24-hours post-injury, etc.) occupies a unique corner of pose space and can be defined by unique sequences of sub-second movements. While meloxicam can relieve mechanical and thermal sensory-reflexive responses, its effect on spontaneous behavior is more subtle as mice injected with either saline or meloxicam at 24-hours post-injury still share common sub-second movements and movement sequences. Thus, our pose estimation results demonstrate that current analgesic treatments for pain are unlikely to revert an animal back to a pre-injury behavioral state. However, the application of combined approaches that can more accurately correlate unique and distinguishable behavior signatures with their underlying pathological mechanisms will drastically improve the translational potential of analgesic drug development from bench-to-bedside.

## Materials and Methods

### Animals (Rutgers University)

Male mice of C57BL/6N background were used for behavioral analyses. All procedures were approved by the Rutgers University Institutional Animal Care and Use Committee (IACUC; protocol #: 201702589). C57BL6 mice were purchased from Jackson Laboratories. All animals were habituated to our facility for 2 weeks after delivery before beginning behavioral experiments described below. All mice used in experiments were housed in a regular light cycle room (lights on from 08:00 to 20:00) with food and water available ad libitum. All cages were provided with nestlets to provide enrichment. All mice were adults between 2 and 4 months. Animals were co-housed with 4 mice per cage in a large holding room containing approximately 300 cages of mice. 20 µl 3% (w/v) λ-Carrageenan (Sigma-Aldrich) in PBS 1X was injected into the mouse left hind paw using a Hamilton syringe. All animals were acclimated to the testing room for an hour prior to testing. For 3D pose imaging, mice were gently placed in the middle of a circular 17” diameter enclosure with 15”-high walls (US Plastics) and allowed to roam freely for 20 minutes while being recorded with the Kinect2 depth-sensing camera. Mice were habituated to the Hargreaves testing chambers for two hours over two days. Thermal hyperalgesia was assessed at baseline and 6- and 26-hours post-carrageenan injection after 3D pose imaging at baseline, 4- and 24-hours post-carrageenan injection. Inflammation was induced while mice were under inhalation anesthesia (2 to 3.5% isoflurane according to mice’s loss of consciousness and anesthetic depth (monitoring of respiratory rate and pattern and responsiveness to toe pinch). Saline or meloxicam (5mg/kg, Henry Schein Animal Health) was injected intraperitoneally 22 hours post carrageenan injection (2 hours before the last 3D pose imaging session).

### Animals (University of Cambridge)

Experiments performed in Cambridge UK (dynamic weight bearing, electrophysiology and immunohistochemistry) were regulated under the Animals (Scientific Procedures) Act 1986 Amendment Regulations 2012. The University of Cambridge Animal Welfare and Ethical Review Body also approved all animal experiments. Cutaneous afferents innervating both hind paws of male C57BL/6J mice were labelled with the retrograde tracer Fast Blue (2% w/v in sterile PBS; Polysciences) when animals were aged 9 weeks. 3 × 1 µl injections were made to the lateral, central and medial plantar aspects of each hind paw under anesthesia (intraperitoneal delivery of ketamine, 100 mg/kg and xylazine, 10 mg/kg) as previously described (da Silva Serra et al., 2016). 5-7 days after retrograde labelling, the right hind paw of mice was inflamed by injection of 20 µl 3% (w/v) λ-carrageenan (Sigma-Aldrich), inflammation was induced while mice were under inhalation anesthesia (2% isoflurane). The diameters of each ankle and foot pad were measured with digital callipers before and 4- or 24-hours post inflammation. 5 animals were used for each time point for electrophysiology experiments, a separate cohort of 4 animals were used for immunohistochemistry experiments.

### Animals (University of Pennsylvania)

Mice for behavior testing were maintained in a barrier animal facility in the Carolyn Lynch building at the University of Pennsylvania. The Lynch vivarium is temperature controlled and maintained under a 12-hr light/dark cycle (7 am/7 pm) at 70 degrees Fahrenheit with ad lib access to food (Purina LabDiet 5001) and tap water. The feed compartment on the wire box lid of the cage was kept at a minimum of 1/3 full at all times. All cages were provided with nestlets to provide enrichment. All procedures were conducted according to animal protocols approved by the university Institutional Animal Care and Use Committee (IACUC) and in accordance with the National Institutes of Health (NIH) guidelines. C57BL mice were purchased from Jackson Laboratories. All animals were habituated to our facility for 2 weeks after delivery before beginning behavioral experiments described below. All mice were adults between 2 and 4 months. Animals were co-housed with 4–5 mice per cage in a large holding room containing approximately 500 cages of mice. 20 µl 3% (w/v) λ-Carrageenan (Sigma-Aldrich) in 0.9% sterile NaCl solution (saline) was injected into the mouse hind paw. Mechanical sensitivity was assessed with PAWS at baseline, 4 and 24 hours post carrageenan injection. Inflammation was induced while mice were under inhalation anesthesia (2 to 3.5% isoflurane according to mice’s loss of consciousness and anesthetic depth (monitoring of respiratory rate and pattern and responsiveness to toe pinch). Saline or meloxicam (5mg/kg, Henry Schein Animal Health) was injected intraperitoneally 22 hours post carrageenan injection (2 hours before the last PAWS behavioral testing session).

### 3D pose analysis

Depth data were modelled as previously published (Wiltschko et al., 2015). First, raw depth frames were collected from a Microsoft Kinect, mounted above the arena, using custom acquisition software written in C#. Frames were collected at 30 Hz, and each frame was composed of 512 × 424 pixels, where each pixel contained a 16-bit integer specifying the distance of that pixel from the sensor in mm. After each session, frames were gzip compressed and moved to another computer for offline analysis. The mouse’s center and orientation were found using an ellipse fit. Then, an 80 × 80 pixel box was drawn around the mouse, and the mouse was rotated to face the right hand side. Next, if the tracking model was used, missing pixels were identified by their likelihood according to the Gaussian model. Low-likelihood pixels were treated as missing data and principal components (PCs) are computed using probabilistic PCA (Roweis, 1998; Tipping and Bishop, 1999). Finally, frames were projected onto the first 10 PCs, forming a 10 dimensional time series that described the mouse’s 3D pose trajectory. These data were used to train an autoregressive hidden Markov model (AR HMM) with 3 lags to cluster mouse behavioral dynamics into discrete ‘‘modules,’’ with state number automatically identified through the use of a hierarchical Dirichlet process. Each state was comprised of a vector autoregressive process that captures the evolution of the 10 PCs over time. The model was fit using Gibbs sampling as described in (Wiltschko et al., 2015) using freely available software (https://github.com/mattjj/ pybasicbayes). Model output was insensitive to all but two hyperparameters, which were set using unsupervised techniques for determining the length scales for discrete behaviors as was previously published (Wiltschko et al., 2015).

### 3D pose analysis: Behavioral usage and transition matrix analysis

Module usage was calculated by summing the number of occurrences of each module and dividing by total module usage across a recording session, converting module usage into a percentage. The number of modules used for each analysis was based on the module usage across all sessions within a condition. Transition matrices were calculated by counting the total number of occurrences module A transitions into module B (for all modules).

### 3D pose analysis: Behavioral Linear Discriminant Analysis

Linear Discriminant Analysis (LDA) was performed using the scikit-learn implementation. Individual normalized usage or bigram transition probabilities were fed to the LDA model, including group labels, and fit to either 2 or 3 components using the eigen solver and an empirically found shrinkage value. Results were then plotted with seaborn and matplotlib.

### Learned Embeddings of module Sequences

Each animal was represented as a doc2vec (“document to vector”) embedding [1] using the Gensim software package [2], version 4.0.1, in Python 3. Doc2vec is a standard unsupervised sequence embedding technique used widely in natural language processing. Each animal and each module was first encoded uniquely as a one-hot vector of 150 (80 animals + 70 modules) dimensions. These one hot vectors were mapped by a linear transformation to an *n*-dimensional embedding space. The linear transformation was adapted by stochastic gradient descent using two losses, resulting in two embeddings. Following Le and Mikolov, 2014, the two embeddings for each animal were averaged to produce the final animal representation. Both embeddings were trained with standard procedures which we outline here. The first embedding was learned by 1) randomly sampling an animal and a contiguous length-2*k* sequence of modules emitted by the animal, 2) mapping the one hot vectors of the sampled animal and modules to the *n*-dimensional embedding space, 3) averaging the animal and module embeddings, 4) mapping the average to the 70-dimensional space of probability distributions over modules, and 5) predicting from this output the identity of a random module in the size-2*k* window. The embedding space was adapted to improve accuracy on this module prediction task. In the parlance of doc2vec methods, this is “distributed memory” (DM) embedding. The second embedding was learned using only representations of animals (no module embeddings). Similar to the first technique, an animal was randomly sampled together with a 2*k*-module subsequence in its raw behavioral sequence. The one hot vector of only the animal was mapped to the embedding space. The animal embedding was then mapped to the 2*k* × 70 dimensional space of distributions on 2*k*-long module subsequences, from which random modules in the sampled subsequence window were predicted. The embedding space was adapted to improve the accuracy of this subsequence prediction task, which also helps the embedding encode higher order information beyond neighboring modules in the raw sequence. This is referred to as a “distributed bag of words” (DBOW) embedding. Again, the final embedding for each animal to be used in classification was the concatenation of DM and DBOW embeddings. In order to determine the best possible embedding model, we performed a hyperparameter grid search over the embedding dimension, (*n*, between 70 and 4900 in 8 logarithmically-spaced steps), the embedding window size, (*k =* 2, 4, 8, 16, 32), the number of training epochs, (*e*=50, 100, 150, 200, 250) and the type of raw data used. For raw data, we either used the modules assigned to every video frame (“frames” data), or the discrete sequence of modules independent of their real-time duration (“emissions” data). The search revealed that the model with the best generalization accuracy (see below) had *n* = 70, *k=4*, *e*=250 and used the emissions data.

### Classifier Analysis of Animal Representations

We used a machine learning classifier to compare the expressive power of three types of animal representations: module usages, module transition probabilities, and learned embeddings. Our goal here was to predict which of the five experimental manipulations an animal received from each of these representations. The dimensionality of the input data was generally different in the three cases, with usages fixed at 70 dimensions. For transitions, we used the most frequent transitions on average, where m was chosen by a grid search having the same values as the n search discussed above (m=1455 was optimal). We trained an identical, L2 -regularized, 5-class logistic regression classifier on each type of representation using leave-one-out cross validation. The regularizer weight was chosen by a grid search over 11 values logarithmically spaced between 1 × 10-5 and 1 × 105 and all results represent the best regularization weight for each model. The classifier was trained using scikit-learn in Python 3. The classifier was trained with a stopping criterion of 1 × 10-5 and with balanced class correction. In this setting, the average validation accuracy for the best models trained on usages, transitions and embeddings were 71.25% 81.25% and 90.00%, respectively. Since the embeddings take into account much larger scale information (i.e. windows of 2k = 8 modules), we can hypothesize consequently that there are experimental effects which differentiate the groups and manifest at behavioral scales beyond usages and one-step transitions.

### Finding characteristic module sequences

There are several possible ways to associate to each embedding a set of characteristic modules or module sequences. For the present study, we adapted a standard method (Mikolov et al., 2013) for scoring the significance of *n*-grams in written text. An *n*-gram is a contiguous sequence of *n* words or, in our case, modules. The original method scored the significance of a bigram *(w1, w2)* by computing

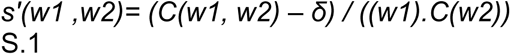

where *C(x)* represents the counts of × in the data and δ controls how many counts are required to make s positive. The idea is to discount the significance of the bigram *(w1 , w2)* by the amount that *w1* and *w2* tend to appear individually (including in the bigram of interest). The original authors detected higher order n-grams by applying this scoring function to sequences repeatedly, thresholding *s* to determine which bigrams should become a fixed phrase, and congealing detected phrases into single symbols.

We took a similar approach with a new emphasis on module n-grams which distinguished between animal classes. To that end, let *A* be the module sequences associated to a given experimental class (e.g. baseline), and *-A* be the complementary module sequences of all other animals in all other classes. Further, for a given *n*-gram *w* = *(w1, …, wn)*, denote the set of contiguous subgrams of lengths 1, …, *n* by S(*w*) = {(*w1*), (*w2*), …,(*w1,w2, …*), (*w2, w3*), …}. Then, we define

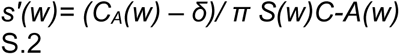

In words, we discount the n-grams detected in class A by the amount that all subgrams, including the n-gram itself, appear in -A. If any subgram was found in A but not in the complement, we gave it a count of ε = 1 × 10-3. δ was set to 1. We collected scores for 2,3 and 4 grams and then found the unique n-grams associated to each class. We also experimented with setting the complement (i.e. -A) as the whole data set (including A), but this produced worse quantitative results than Eq. S.2.

To show that these n-grams have a material effect on the quality of the learned embeddings, we then ablated them and replaced them with random modules in the raw data. The ablations of the five data sets (baseline, 4 hours carrageenan, 24 hours saline, 24 hours meloxicam and baseline meloxicam) totaled approximately 0.10%, 3.28%, 0.09%, 0.11%, 0.08% of the raw modules. Averaging over ten realizations of random module insertions, accuracy dropped from 90% to 83.25 % +/- 1.70 % (s.e.). Since the optimal window size was 2k = 8, performance would likely have degraded even more had we ablated higher order n-grams, though these are difficult to find. Note that performance in the random control condition still degrades slightly, because informative features were randomly ablated. The top ranked n-grams for each class are given in Table S.1. Scores varied over a very large range (4.68 × 10-2 to 8.56 × 105), with the highest scores arising from n-grams for which many subgrams had to be replaced by ε. This naturally occurs for longer n-grams.

### Behavior: Hargreaves Assay

To assess hind paw heat sensitivity, Hargreaves’ test was conducted using a plantar test device (IITC). Mice were placed individually into Plexiglas chambers on an elevated glass platform and allowed to acclimate for at least 30 minutes before testing. A mobile radiant heat source of constant intensity was then applied to the glabrous surface of the paw through the glass plate and the latency to paw withdrawal measured. Paw withdrawal latency is reported as the mean of three measurements for both hind paws with at least a 5 min pause between measurements. A cut-off of 20 s was applied to avoid tissue damage.

### Behavior: Dynamic Weight Bearing

The weight bearing of free-moving animals was assessed using a dynamic weight bearing apparatus (Bioseb). Each test lasted three minutes, mice were naïve to the test device before baseline weight bearing was assessed. The 2 highest confidence levels of automatic paw assignment by the accompanying software were taken forward for analyses; correct paw assignment was manually validated for at least 1 minute 30 seconds of each test.

### Electrophysiology

Mice were sacrificed by cervical dislocation before the lumbar DRG (L2 – L5) from carrageenan injected and non-injected sides were collected separately in dissociation media (L-15 + GlutaMAX growth media supplemented with 24 mM NaHCO_3_; Life Technologies). Dissected DRG were then incubated in dissociation media containing 1 mg/ml type 1A collagenase (Sigma Aldrich) and 6 mg/ml bovine serum albumin (BSA; Sigma-Aldrich) for 15 min at 37 °C, 5% CO_2_, before a further 30 min in dissociation media containing 1 mg/ml trypsin (Sigma-Aldrich) and 6 mg/ml BSA. DRG were then suspended in culture media (L-15 + GlutaMAX growth media supplemented with 10% (v/v) fetal bovine serum, 24 mM NaHCO_3_ 38 mM glucose and 2% (v/v) penicillin/streptomycin) before several rounds of mechanical trituration and brief centrifugation (160g, 30 s). After sufficient trituration, dissociated cells were pelleted (160g, 5 min), resuspended in culture media and plated on poly-D-lysine/laminin coated glass coverslips (BD Biosciences) and incubated at 37 °C, 5% CO_2_. Electrophysiology experiments were performed the following day, recordings were made using an EPC-10 amplifier (HEKA) and corresponding Patchmaster software. The extracellular solution contained (in mM): NaCl (140), KCl (4), MgCl_2_ (1), CaCl_2_ (2), glucose (4) and HEPES (10), pH 7.40. Patch pipettes were pulled from borosilicate glass capillaries (Hilgenberg) using a P-97 pipette puller (Sutter Instruments) with resistances of 4-8 MΩ and back filled with intracellular solution containing (in mM): KCl (110), NaCl (10), MgCl_2_ (1), EGTA (1), HEPES (10), Na_2_ATP (2), Na_2_GTP (0.5), pH 7.30. Whole cell currents or voltages were sampled at 20 kHz from Fast Blue labelled neurons, identified by LED excitation at 365 nm (Cairn Research). Step wise depolarization (Δ10 pA, 50 ms) was used to determine the action potential threshold of cells. Only cells which fired action potentials and had a resting membrane potential less than or equal to -40 mV and were included in analyses. Action potential parameters were measured using Fitmaster software (HEKA) and IgorPro software (Wavemetrics) as previously described (Chakrabarti et al., 2018). The excitability of neurons was further assessed by applying a suprathreshold (2x action potential threshold) for 500 ms, the frequency of action potentials during this time was noted. The activity of macroscopic voltage-sensitive channels was assessed in voltage clamp mode with appropriate compensation for series resistance. Cells were held at -120 mV for 150 ms before stepping to the test potential (−60 mV – 55 mV in 5 mV increments) for 40 ms and returning to a holding potential of -60 mV for 200 ms between steps. Peak inward and outward currents were normalized to cell size by dividing by cell capacitance. Peak inward current densities were then fit to a Boltzmann function to determine the reversal potential and half-activating potential of voltage sensitive channels. The conductance of channels guarding inward currents was determined using the equation, G = I/(V_m_-E_rev_), where G, conductance; I, peak inward current; V_m_, mV step to elicit I and E_rev_, reversal potential and fit to a Boltzmann function. To compare macroscopic voltage-sensitive currents between neurons isolated from the inflamed and non-inflamed sides peak current densities and conductance values were normalized to those obtained from cells from the contralateral side.

### Immunohistochemistry

24-hours post inflammation a cohort of 4 mice were transcardially perfused with 4% (w/v) PFA under terminal anesthesia (intraperitoneal delivery of 200 mg/kg sodium pentobarbital). Lumbar DRG (L3 – L4) were then collected from both the inflamed and non-injected sides and post-fixed in Zamboni’s fixative for 30 min, followed by overnight incubation in 30% (w/v) sucrose at 4 °C for cryoprotection. Individual DRG were then snap-frozen in Shandon M-1 Embedding Matrix (Thermo Fisher Scientific). 12 µm sections of each DRG were collected on a cryostat sequentially across 10 slides. After washing with PBS containing 0.001% (v/v) Tween-20 (Thermo Fisher Scientific), slides were incubated with antibody dilutant (1 % (w/v) BSA, 5% (v/v) donkey serum and 0.02% (v/v) Triton-X-100 in PBS) at room temperature for 1 hour before overnight incubation at 4 °C with an anti-TRPV1 antibody (1:500; guinea-pig polyclonal; Alomone, AGP-118). The following day slides were washed three times with PBS-Tween before incubation with donkey anti guinea-pig IgG-AF488 for 2 hours at room temperature, followed by a further two washes and mounting. Images were acquired with an Olympus BX51 microscope and Q-Imaging camera. Two sections per DRG per animal were analyzed, briefly, each cell was selected as a region of interest and individual cells were considered positively stained if the background corrected intensity exceeded 2x SD of the normalized intensity across all sections. Negative controls which were not exposed to any primary antibody showed no fluorescent staining.

### Data Analysis

Data are presented as mean ± standard error of the mean (SEM) for cellular experiments or mean ± standard deviation (SD) for *in vivo* experiments. Statistical tests used to assess differences between groups are detailed in individual figure legends. Behavioral assays were replicated several times (3 to 10 times depending on the experiments) and averaged per animal. Statistics were then performed over the mean of animals. Statistical analysis was performed in GraphPad Prism (USA) using two-sided paired or unpaired Student’s t-tests, one- or two-way repeated-measures ANOVA for functional assessments, when data were distributed normally. Post hoc Tukey’s or Bonferroni test was applied when appropriate. The significance level was set as p < 0.05. The nonparametric Mann-Whitney or Wilcoxon signed rank tests were used in comparisons of <5 mice.

### High-speed imaging and video storage

Mouse behaviors were recorded at 2000 fps with a high-speed camera (Photron FastCAM Mini AX 50 170 K-M-32GB - Monochrome 170K with 32 GB memory) and attached lens (Zeiss 2/100M ZF.2-mount). Mice performed behavior in rectangular plexiglass chambers on an elevated mesh platform. The camera was placed at a ∼45° angle at ∼1–2 feet away from the Plexiglas holding chambers on a tripod with geared head for Photron AX 50. CMVision IP65 infrared lights that mice cannot detect were used to adequately illuminate the paw for subsequent tracking in ProAnalyst. All data were collected on a Dell laptop computer with Photron FastCAM Analysis software (average size of video file = ∼2 GB).

### Somatosensory behavior assays

In all behavioral experiments, we used a sample size of 6–10 mice per strain, as these numbers are consistent with studies of this kind in the literature to reach statistically significant conclusions. All mice were habituated for 2 days, for one hour each day, in the Plexiglas holding chambers before testing commenced. Mice were tested in groups of five and chambers were placed in a row with barriers preventing mice from seeing each other. On testing day, mice were habituated for an additional ∼10 min before stimulation and tested one at a time. Stimuli were applied through the mesh to the hind paw proximal to the camera. Testing only occurred when the camera’s view of the paw was unobstructed. Mice received two stimuli on a given testing day (db and lp) and were given at least 24 hr between each stimulus session. Stimuli were tested from least painful to most: dynamic brush then light pinprick. Dynamic brush tests were performed by wiping a concealer makeup brush (L’Oréal Paris Infallible Concealer Brush, item model number 3760228170158) across the hind paw from back to front. Light pinprick tests were performed by touching a pin (Austerlitz Insect Pins) to the hind paw of the mouse. The pin was withdrawn as soon as contact was observed.

### Automated paw tracking

#### Proanalyst

We used ProAnalyst software to automatically track hind paw movements following stimulus application. This software allowed us to integrate automated and manually scored data, possible through the ‘interpolation’ feature within ProAnalyst. We were able to define specific regions of interest (paw), track, and generate data containing ‘x’ and ‘y’ coordinates of the paw through time. In a subset of videos, additional manual annotation was performed for increased accuracy.

#### DeepLabCut

For deep learning-based paw tracking in DeepLabCut (DLC), we pseudo-randomly selected a subset of training frames from trials which contained the greatest behavioral variation and hand-labeled the hind paw toes, center, and heel. We trained DLC to predict toe, center, and heel positions in unlabeled video frames.

### Quantifying withdrawal behavior

#### PAWS

Behavioral features were extracted from raw paw position time series in an automated and standardized procedure. First, the start and end of paw movement (paw at rest on the ground) were identified, and analysis was restricted to this time window. Peaks in paw height were then determined based on Savitsky-Golay smoothed estimates of paw velocity, and the first peak identified. The time of the first peak (designated t*) was used to separate pre-peak behavioral feature calculations from post-peak calculations. To differentiate shaking from guarding in the post-peak period, we constructed a moving reference frame based on the principal axis of paw displacement across a sliding window (0.04 s in duration) for each time point, and identified periods of consecutive displacements above a specified threshold (35% of maximum paw height) as periods of shaking. Note that in the construction of the moving reference frame the principal axes of variation were recovered via principal component analyses, which is not invariant to the sign of the recovered axes. Since displacement is measured over time it is sensitive to reversals in sign along the axis we measure it. We therefore ensured consistency by using the axis direction minimizing the angular deviation from the axis recovered at the previous time step. PAWS is open source and freely available at https://github.com/crtwomey/paws.

#### B-SOiD

To determine behavioral sub-actions following foot stimulation, pose estimation data was passed along to the unsupervised behavioral discovery and extraction algorithm, B-SOiD. Experimentalists processing this data were blind to the experimental condition. Position coordinates of fore paw hind paw motion tracked by DeepLabCut in samples of 2000fps video, and then were imported to the B-SOiD app to identify unique behavioral clusters in response to pain stimulus. Data from dynamic brush and light prick Hour-4 sessions were combined to develop a generalized B-SOiD model of pain response. The frame rate was scaled down to 1/7th of the original to help B-SOiD extract sub behavioral features. Data used in the B-SOiD model were the hind paw toe, hind paw center-paw and two static reference positions. The two static reference points were the initial positions of the fore paw the maximum elevation of toe post stimulation. B-SOiD performed nonlinear embedding to transform 16-dimensional data to 5-dimensional UMAP space. The 16-dimensional data include frame by frame calculation of distance and angle between all four points as well as the speed of the two body parts. 11 behavioral clusters were identified and used to train the random forest classifier of the algorithm, which was then used to assign behavioral labels to data from all epochs and stimulation types. Every frame was labeled, and a smoothing kernel was used to eliminate any sub actions lasting under 2.5 seconds (5 frames). Importantly, beyond the spatiotemporal relationship values between the points, no other information was available to the algorithm. B-SOiD is open source and freely available at https://github.com/YttriLab/B-SOID.

## Acknowledgements

LAP and EStJS acknowledge the work of University of Cambridge Combined Animal Facility staff. LAP was supported by the University of Cambridge BBSRC Doctoral Training Programme (BB/M011194/1). EStJS acknowledges support from Versus Arthritis (RG 21973). IAS, WF, HR, JA, ZAJ were supported from startup funds from the University of Pennsylvania and Columbia University, a grant from the National Institutes of Health NIH/NIDCR, R00-DE026807, the Rita Allen Foundation, and the Alfred P. Sloan Foundation. VEA, MAT, MB are supported by startup funds from Rutgers University. VEA and MB are also supported by the Pew Charitable Trust. MAT is also supported by Tourette Association of America, New Jersey Center for Tourette Syndrome, and a Robert Wood Johnson foundation grant.

## Author Contributions

LAP and MB contributed equally to the work. EStJS, IAS, and VEA conceived the research study. EStJS, IAS, EY, and VEA designed experiments. LAP performed the electrophysiology experiments. MB, HR, ZAJ, and WF performed the behavioral experiments. ZAJ, WF, JA, NM, and EY performed the analysis for PAWS and B-SOiD. MB, MAT, JKT, MR performed the analysis for Motion Sequencing. MB, LAP, IAS, HR, ZAJ, WF, EStJS, MR, and VEA wrote the manuscript. All authors contributed to the editing of the final document.

## Inclusion and Diversity Statement

All the authors in this study actively support inclusion, diversity, and equality in science. The two senior authors, and several co-authors are from racial and ethnic groups underrepresented in science. Gender and sexual orientation diversity is also highlighted in the composition of this team, including representation of women as the first and last author. One or more of the authors of this paper received support from a program designed to increase minority representation in science, including the PDEP fellowship from Burroughs Wellcome Fund. While citing references scientifically relevant for this work, we also actively worked to promote gender balance in our reference list.

## Competing Interests

The authors declare no conflict of interest with any of the data presented in this article.

---

**Table S1.**
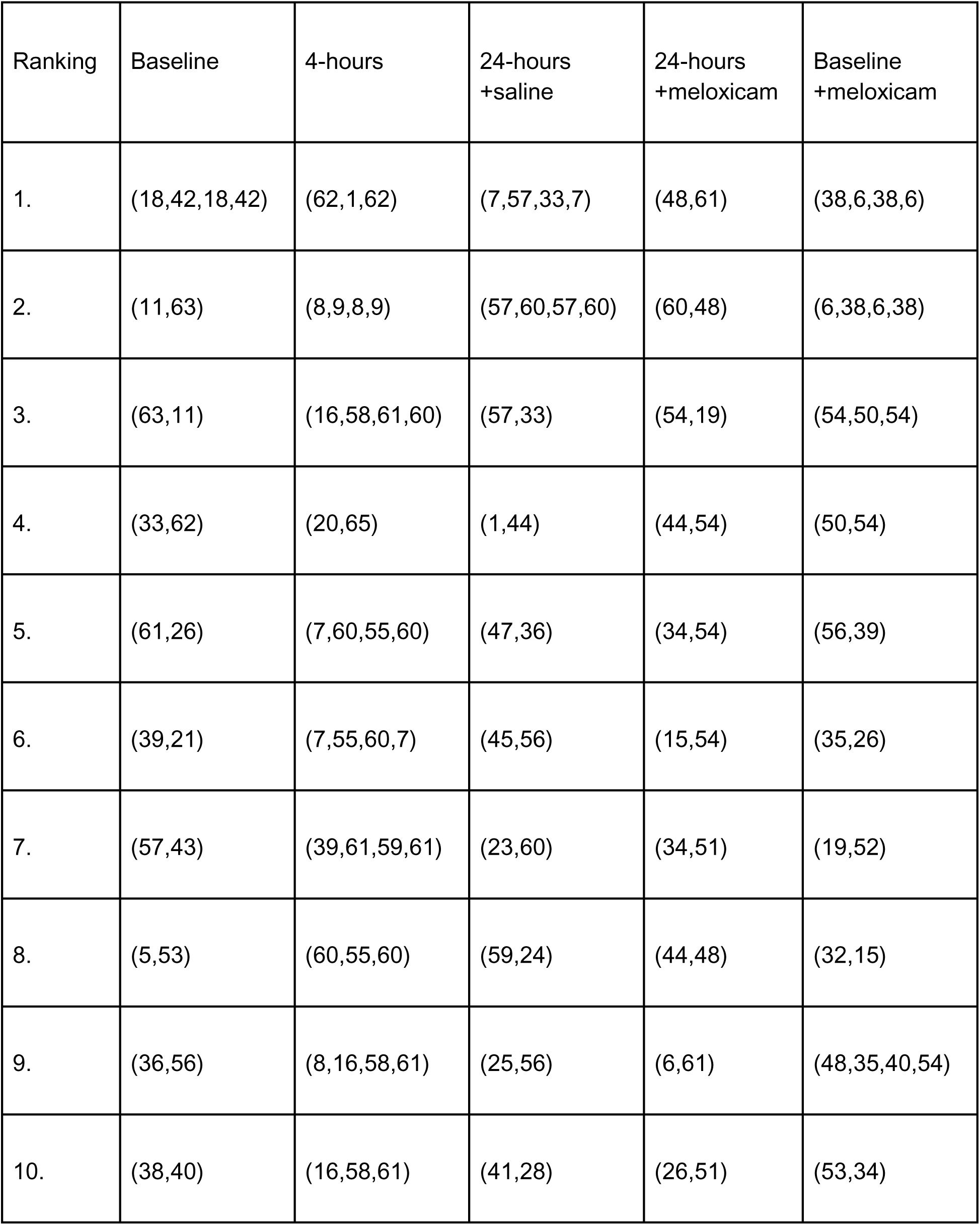
Top ranked module sequences for each behavioral state.

| Ranking | Baseline | 4-hours | 24-hours<br>+saline | 24-hours<br>+meloxicam | Baseline<br>+meloxicam |
| --- | --- | --- | --- | --- | --- |
| 1. | (18,42,18,42) | (62,1,62) | (7,57,33,7) | (48,61) | (38,6,38,6) |
| 2. | (11,63) | (8,9,8,9) | (57,60,57,60) | (60,48) | (6,38,6,38) |
| 3. | (63,11) | (16,58,61,60) | (57,33) | (54,19) | (54,50,54) |
| 4. | (33,62) | (20,65) | (1,44) | (44,54) | (50,54) |
| 5. | (61,26) | (7,60,55,60) | (47,36) | (34,54) | (56,39) |
| 6. | (39,21) | (7,55,60,7) | (45,56) | (15,54) | (35,26) |
| 7. | (57,43) | (39,61,59,61) | (23,60) | (34,51) | (19,52) |
| 8. | (5,53) | (60,55,60) | (59,24) | (44,48) | (32,15) |
| 9. | (36,56) | (8,16,58,61) | (25,56) | (6,61) | (48,35,40,54) |
| 10. | (38,40) | (16,58,61) | (41,28) | (26,51) | (53,34) |

**Extended Data Figure 1.**
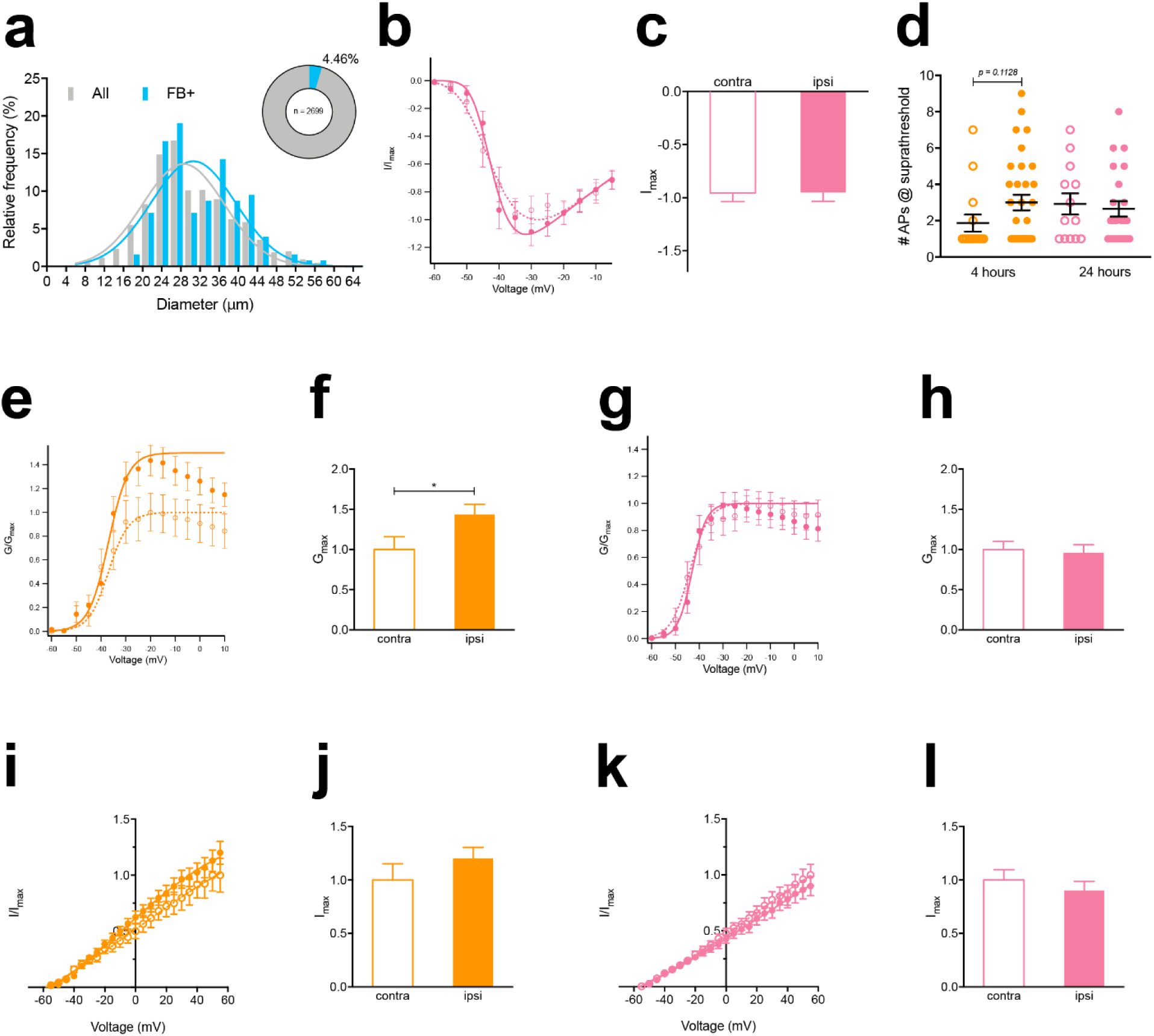
Unilateral injection of carrageenan into the hind paw of mice results in swelling, pain-like behaviors. **(a)** Relative frequency distributions of dissociated sensory neuron soma diameter from hind paw innervating and total lumbar populations, Insert: Proportion Fast Blue positive cells from acutely dissociated cultures of lumbar DRG. Inward macroscopic current densities normalized to the average peak inward current density of contralateral cells **(b)** 24-hours post inflammation and **(c)** Normalized peak inward current densities. **(d)** Number of action potentials (APs) discharged following stimulation of sensory neurons with a suprathreshold (2X rheobase). **(e,g)** Calculated conductance of inward voltage-gated currents normalized to average peak conductance of contralateral neurons. **(f,h)** Normalized peak conductance of inward voltage-gated currents. Outward macroscopic current densities normalized to the average peak outward current density of contralateral neurons either **(i)** 4- or **(k)** 24-hours post inflammation. **(j,l)** Normalized peak outward current densities. * p < 0.05, unpaired t-test.

**Extended Data Figure 2.**
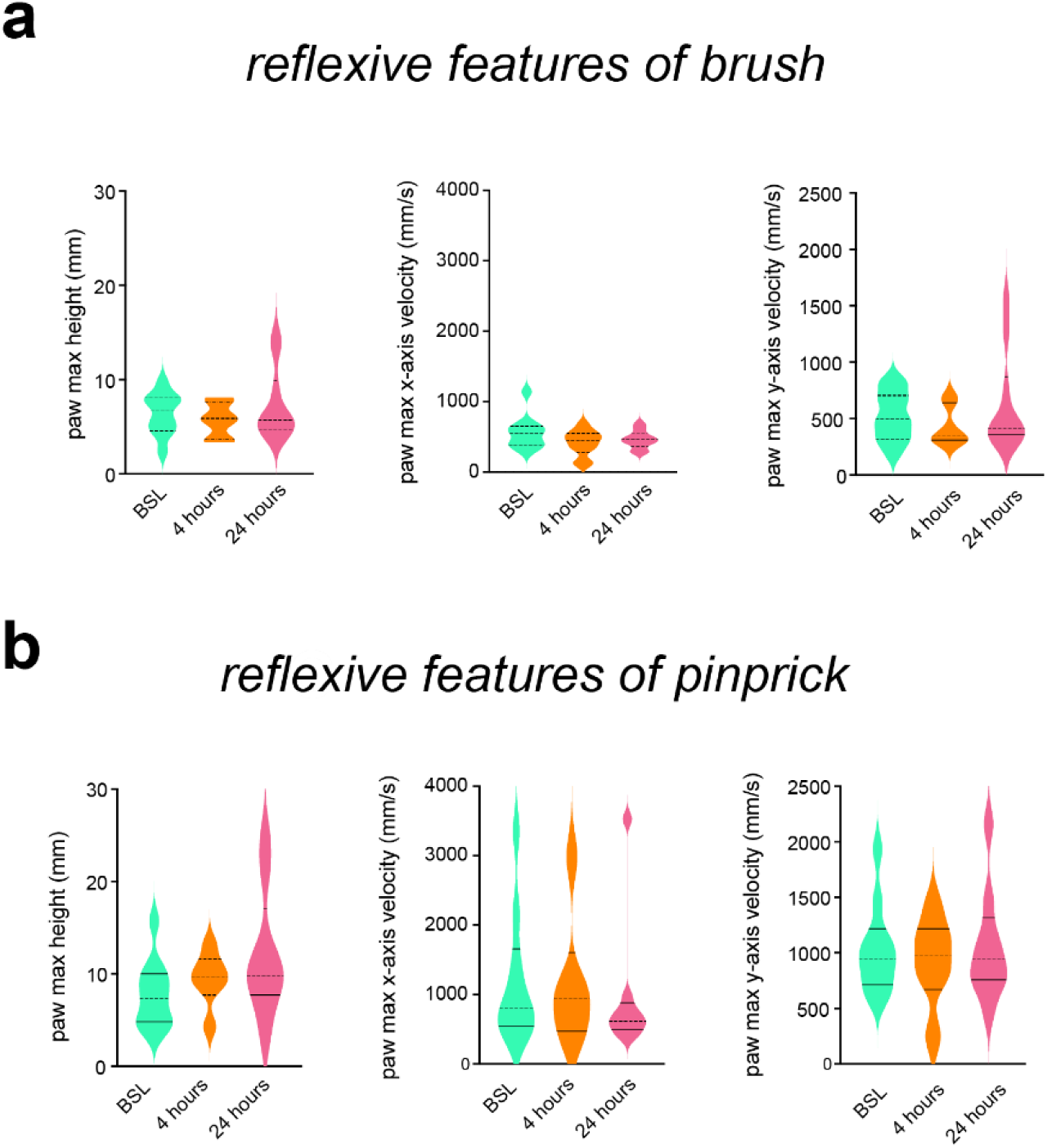
PAWS and B-SOiD automated pain assessment platforms detect defensive coping behaviors associated with pain sensation during inflammation. Reflexive features of the behavioral response to a stimulation with brush **(a)** or pinprick **(b)** at baseline, 4- and 24-hours post-carrageenan injection. Reflexive features do not appear deregulated following induction of inflammation with carrageenan.

**Extended Data Figure 3.**
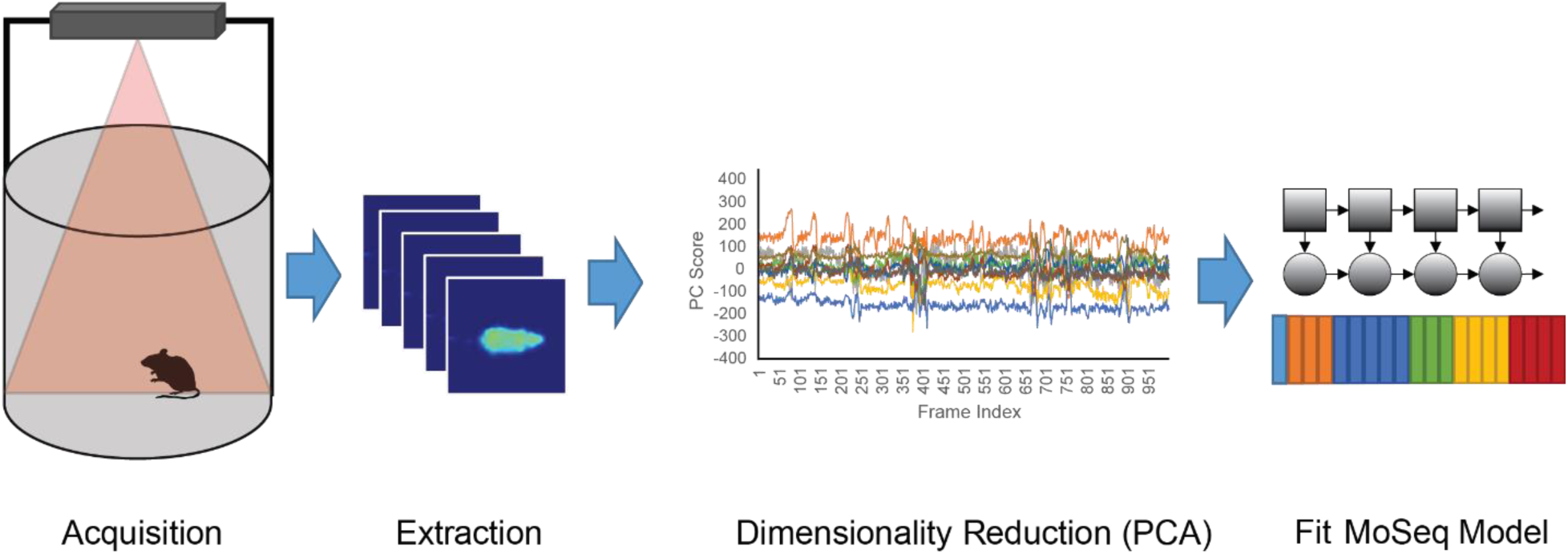
3D pose analysis pipeline. 3D pose analysis uses time-of-flight infrared cameras to detect mouse body contours, depth and movement during 20-min long sessions. After extraction of the frames, reduction of dimensionality by Principal Component Analysis (PCA) followed by fitting an auto-regressive semi-hidden Markov Model (AR-HMM) to our data allows us to identify the set of milliseconds long movement (a.k.a. “modules”) that best describe mouse behavior.

**Extended Data Figure 4.**
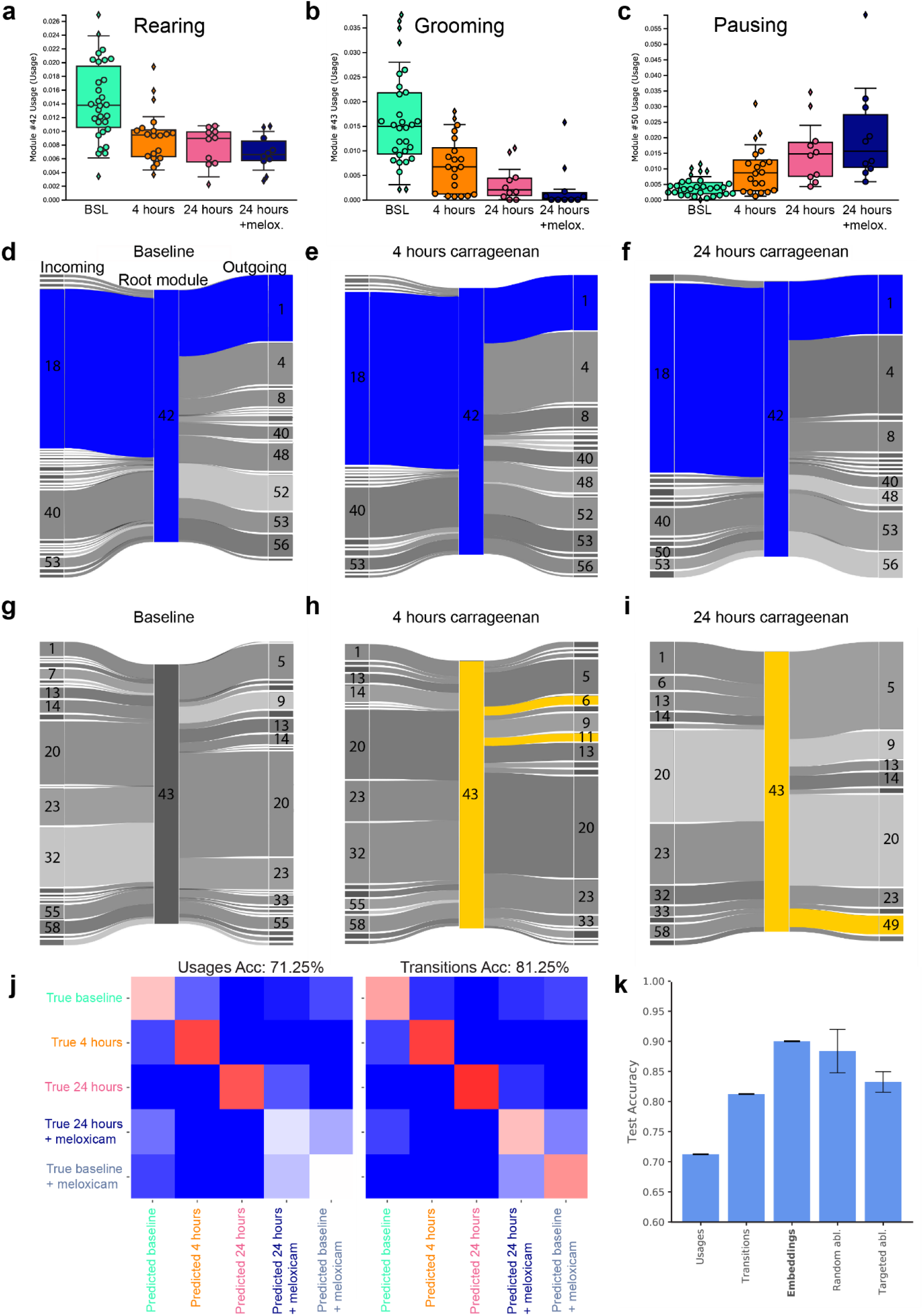
Context-dependent learned embeddings are more predictive of experimental groups than raw usage or transition probabilities and show that meloxicam does not promote return to pre-inflammation spontaneous behavior. **(a,b,c)** Example of modules with varying usages at baseline, and following carrageenan injection at 4-hours, 24-hours and 24-hours with meloxicam injection. **(a)** Example of a rearing module (42, rear against the wall) significantly less used at 24- vs 24-hours+meloxicam. **(b)** Example of a grooming module (43). **(c)** Example of a pause module (50, pause with head up/observe). Overall, we find that intraperitoneal injection of the anti-inflammatory drug meloxicam, which relieves evoked pain, seems to exacerbate the change in spontaneous behavior observed when animals develop inflammatory pain. **(d,e,f,g,h,i)** Sankey diagram representation of 3-long module sequences based on root modules 42 **(d,e,f)** and 43 **(g,h,i)** and their incoming and outgoing modules at baseline, 4- and 24-hours after carrageenan injection. Modules 42 and 43 represent examples of rearing and grooming behavior respectively, which usage is downregulated in pain states. Despite decreased usage, we observe a majority of conserved incoming and outgoing modules (green for module 42) **(d,e,f)**. However, we notice the appearance of unique sequences in pain states absent at baseline, such as 43→6 or 43→11 at 4-hours, or 43→49 at 24-hours (yellow for module 43) **(h,i)**. **(j)** Held-out data confusion matrices for an identical classifier trained on raw usage data and transition probabilities with leave-one-out cross validation. Usages and transition probabilities have lower prediction accuracy (71.25% and 81.25% respectively) than context-dependent learned embeddings (See Figure 4). **(k)** Bar plot showing the relative performances of the different representations along with the performance of targeted vs random ablations (abl. = ablation, Acc = accuracy).

## Notes

### Competing Interest Statement

The authors have declared no competing interest.

